# SCI-VCF: A cross-platform GUI solution to Summarise, Compare, Inspect, and Visualise the Variant Call Format

**DOI:** 10.1101/2023.08.09.552664

**Authors:** Venkatesh Kamaraj, Himanshu Sinha

## Abstract

As genomics advances swiftly and its applications extend to diverse fields, bioinformatics tools must enable researchers and clinicians to work with genomic data irrespective of their programming expertise. We developed SCI-VCF, a Shiny-based comprehensive analysis utility to summarise, compare, inspect, analyse, and design interactive visualisations of the genetic variants from the variant call format. With an intuitive GUI, SCI-VCF aims to bridge the approachability gap in genomics that arises from the existing predominantly command-line utilities. SCI-VCF is written with R and is freely available at https://github.com/HimanshuLab/SCI-VCF. For installation-free access, users can avail of an online version at https://ibse.shinyapps.io/sci-vcf-online/.

## INTRODUCTION

Genomics is a rapidly evolving field that has profoundly impacted biomedical and public health research. With the advent of next-generation sequencing (NGS), an increasing number of genomes from various species are getting sequenced, enabling the applications of genomics to proliferate into diverse disciplines such as medicine (Hudson, 2011), agriculture (Wang et al., 2017), microbiology (Padmanabhan et al., 2013), ecology (Klaper and Thomas, 2004), evolutionary biology (Rokas and Abbot, 2009), psychology (Hoehe and Morris-Rosendahl, 2018), and anthropology (Benn Torres, 2020). This is accelerated further by combining genomics with advancements in data science and artificial intelligence (Navarro et al., 2019). Such interoperability is facilitated by well-defined data formats that represent diverse biological entities. One such data structure is the Variant Call Format (VCF).

VCF is a standard file format to store the sequence variations in a genome like single nucleotide polymorphisms (SNP), insertions, deletions (INDEL), structural variants (SV), and other assorted variants (Danecek et al., 2011). It is a tab-delimited text file, with each row comprising information about a genomic variant site and metadata of these variants presented at the beginning of the file. The first eight columns of the headers are fixed and are named CHROM, POS, ID, REF, ALT, QUAL, FILTER, and INFO. Figure 1A depicts the structure and contents of a generic VCF file. It is a widely adopted file format used in many bioinformatics tools that analyse genomic data. While the VCF has many advantages for storing and sharing genomic variants efficiently, it can be complex and arduous to understand, particularly for those unaccustomed to bioinformatics and genomic data analysis.

**Figure 1:**
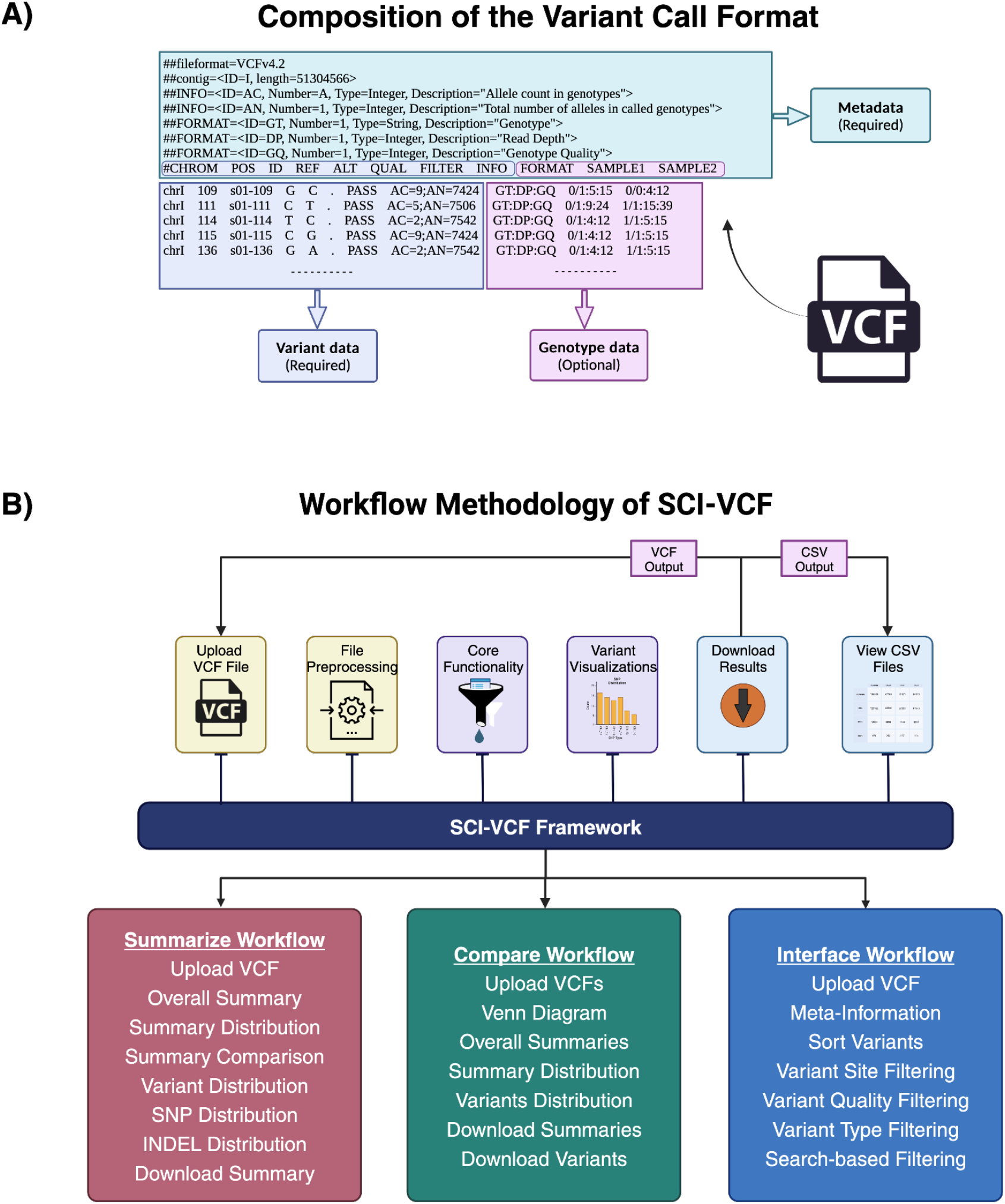
VCF analysis workflow in SCI-VCF. **A)** Composition of the standard VCF. Metadata and the information of variants in the first eight columns are mandatory. Genotype information is optional and can be present across multiple columns. **B)** Workflow methodology of SCI-VCF describing the core functionalities offered by the tool. Created with Biorender.com

Several tools allow users to examine, analyse and interpret VCF files. However, they are predominantly command-line utilities and require significant programming expertise from the user. Data visualisation is a ubiquitous method for quickly understanding complex information, but tools that visualise a VCF file are scarce. VIVA (Tollefson et al., 2019), vcfR (Knaus and Grünwald, 2017), and the vcflib (Garrison et al., 2022) are some existing frameworks that allow the visualisation of VCF. However, effectively using these variant visualisation tools also requires proficiency in programming on the user’s end.

As the applications of genomics burgeon into diverse fields, it calls for tools and software that empower clinicians and researchers to work with genomic data formats irrespective of their programming expertise. Here, we introduce SCI-VCF, a comprehensive toolkit with an intuitive Graphical User Interface (GUI) that lets users summarise, interpret, and compare genomic variants from VCF files. It also equips users to design interactive visualisations of the VCF in numerous ways. SCI-VCF is platform-agnostic and works seamlessly across any operating system. A web version of the tool is also made available for enhanced accessibility. While it is infeasible for a single GUI-based tool to support the wide-spanning analytic capabilities of the VCF, SCI-VCF provides a well-founded framework that simplifies the core components of VCF analyses, thus increasing the approachability of genomics to novices.

## METHODS

SCI-VCF is written in R, a free and open-source programming language for statistical computing, data analysis, and visualisation. The user interface and the routines are designed and developed in Shiny, an R framework to build interactive applications. VCF files are parsed in SCI-VCF using the *vcfR* library and are processed using the *tidyverse* packages. The static visualisations of the VCF files are developed with *ggplot2* (Wickham, 2011), which operates on a grammatical theory of graphics. The static plots are then transformed into interactive visualisations using *Plotly* for R. The area-proportional Venn diagrams are made with the *eulerr* package, and the interactive data tables enabling detailed inspection of VCF files are created using the *reactable* library.

SCI-VCF has three major workflows separated into respective modules in the navigation bar: Summarise, Compare and Interface. Submodules capturing the result of individual analyses are assembled under the major module’s sidebars. Figure 1B elucidates the major workflows of the tool. Both compressed and uncompressed VCF files are accepted as valid inputs in all three major modules. A module to inspect Comma Separated Values (CSV) files is also added to provide a tabular variant-level analysis system. Every plot generated can be customised to equip the user to change various aspects of the visualisation. All the data and visualisations generated by the tool are downloadable for further examination. Sample VCF files are provided to get started with the tool from the ground up.

## RESULTS AND DISCUSSION

### SCI-VCF provides summary statistics of a VCF file in a graphical format

The first four columns of a VCF file can be used to identify the variants present in a normalised VCF file uniquely. Normalising is achieved by breaking down comma-separated multiallelic sites into individual entries and removing duplicates. The nature of REF and ALT entries enabled the classification of variants into different variant types: SNPs, transitions, transversions, INDELs, insertions and deletions individually, and other Multi Nucleotide Polymorphisms, multi-allelic sites and assorted variants that do not fit the other categories. The summary of a VCF file was generated by classifying variants and summing up unique entries in each category.

Statistics derived from this summary, like the ratio of transitions to transversions, are generally used as quality control metrics to assess the confidence in the variants captured by the variant calling algorithm (Wang et al., 2015). The occurrence of variants throughout the genome is presented as an interactive histogram to recognise regions that are depleted or enriched with genetic variations. Such a plot aids in identifying the highly polymorphic and invariant sections of the genome, possibilities of selection pressures, and genomic loci linked to fundamental biological functions and salient clinical features. With the distribution of INDEL sizes captured by SCI-VCF, it is possible to gauge the prevalence of frameshift mutations and structural variants. The spread of each variant type across different chromosomes is represented in diverse plots. By illustrating the synopsis and distribution of specific variant types across the genome, SCI-VCF helps researchers gain valuable insights into the genetic diversity, evolutionary history, and disease susceptibility of an individual, a population, or a species.

### Comparison of a pair of VCF files to understand genetic diversity

Two VCFs were compared by interpreting the first eight columns of the files as two-dimensional heterogeneous tabular datasets. The area-proportional Venn diagram created assists in a quick overview of the commonalities, dissimilarities, and relationships between the two variant calls. The shared and unique variant sets between the two files are summarised and visualised in multiple ways to advance further investigation by the user. Comparing VCFs and recapitulating the resulting variant sets enables the comprehension of the genetic diversity across individuals and populations and aids in gaining crucial insights about them. For example, novel variants with potential clinical and biological significance could be uncovered by comparing the VCFs of an individual or population with a reference database like the 1000 Genomes project (1KGP) (Auton et al., 2015) or the gnomAD dataset (Karczewski et al., 2020). Variant calling pipelines could be validated for consistency and accuracy by comparing their results with available high-confidence variant sets for thoroughly studied samples like the Genome In A Bottle (GIAB) benchmark sets (Zook et al., 2016).

The genetic basis of various diseases is understood by juxtaposing the variants in individuals with and without a particular disease. This procedure helps in identifying the mutations associated with the condition. A similar approach is also used to study genotype-phenotype association and pin down genetic markers for phenotypic traits. SCI-VCF equips users with the framework to effortlessly compare VCFs and summarise the overlapping and dissimilar variants between individuals and populations. These comparisons and summaries are pivotal for understanding genetics, unravelling disease mechanisms, developing targeted interventions, and advancing genomic research.

### SCI-VCF provides in-depth visualisation of the contents of a VCF file

SCI-VCF offers a framework to view, search, sort, identify, and filter the contents of a VCF file. Entries in a file are filterable in terms of keywords, variant sites, quality scores and variant types. Variant sorting and quality filtering are standard analyses when dealing with VCF files, quality filtering aiding in removing the low-quality variants that might have resulted from sequencing errors or mapping artefacts. The variant site and type filtering enable the prioritisation of variants and help focus the analysis on specific regions of interest, such as genes, exons, or regulatory elements. The keyword search was extended to each VCF column, enabling sophisticated filtration capabilities, including annotation-based variant extraction. These variant inspections can be beneficial when studying variants in candidate genes or genomic regions associated with a particular phenotype or disease.

The meta-information from VCF files is extracted and displayed in a searchable fashion to provide additional context about the variants, like the reference genome used, databases used for variant annotation, descriptions of those annotations, and possible pre-processing steps done with the VCF file. The filtered variants and the meta-information can be downloaded by the user for further study. While the results obtained in the VCF file type can then be summarised and compared using the respective modules in SCI-VCF, the results downloaded in CSV file type can be inspected in-depth in the ‘View CSV Files’ module.

All data visualisations in SCI-VCF are presented as interactive plots to enhance exploratory data analysis with VCF files. With the help of features like input selection, zooming, panning, and tooltips, researchers can explore different aspects and dive deeper into specific areas of interest to uncover patterns, outliers, and relationships that may not be apparent in static plots. All graphics made with the tool are supplemented with plot customisation features to improve the effectiveness, clarity, and visual impact of the data visualisations. As points of interest from the displayed interactive plots can be customised and saved locally, users can extract publication-ready visualisations from SCI-VCF.

### Use case for SCI-VCF application

To demonstrate its utility, we performed a case study only using SCI-VCF. Figure 2 displays the visualisations taken directly from SCI-VCF at various stages of this analysis. The input for the study contained variants captured by the BWA-GATK pipeline in the whole genome sequence (WGS) of the GIAB sample HG002. A total of about 4.78M variants were present in the file. Figure 2A shows the summary of the entries present in the file. Nearly 80% of the variants in the file were SNPs, with a Ts/Tv of 2.02, which was the expected value for the human WGS variants. By examining the INDEL size distribution plot, it was evident that no structural variants were present in the file. This result was anticipated as the variant calling workflow was not explicitly designed to capture structural variants. The maximum size of an insertion was 456 bp, and deletion was 339 bp, with most INDELs biased towards smaller sizes. The functionalities to interact with the visualisations were seamlessly integrated within SCI-VCF, and the same was used to zoom in on the INDEL size distribution plot to limit the sizes to 50 bps on both axes in Figure 2B.

**Figure 2:**
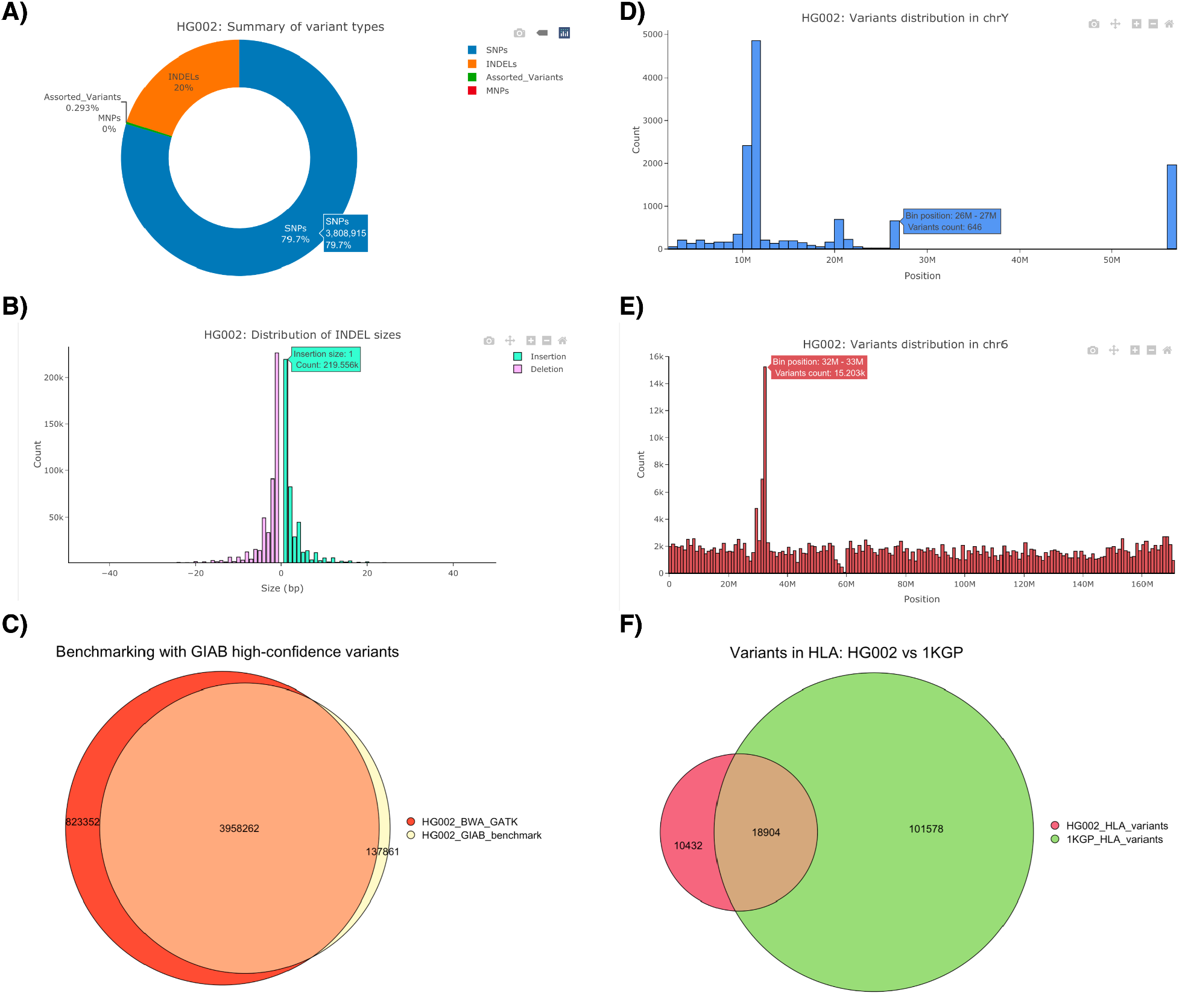
Analyses of variants in HG002 using SCI-VCF. **A)** Distribution of variant types; **B)** Size distribution of INDELs; **C)** Comparison of HG002 variants with the GIAB high-confidence variants; **D)** Distribution of variants in chrY; **E)** Distribution of variants in chr6; **F)** Comparison of variants in the HLA region with the same from the 1000 genomes project.

To validate the variant calling pipeline used to create the VCF file, we benchmarked its output with the high-confidence VCF file provided by the GIAB consortium. For this task, we used the ‘Compare’ workflow and the results in the form of a Venn diagram are depicted in Figure 2C. The pipeline captured nearly 96.6% of the high-confidence variants released by GIAB, ensuring its credibility. Further study of the position-level distribution of variants revealed that chrY had a large genomic region, nearly 30 Mbp, with no variants. Meanwhile, the 32-33 Mbp genomic locus in chr6 contained over 15,000 variants, the maximum observed value for any 1 Mbp genomic window. Figures 2D and 2E depicted the distribution of variants in chrY and chr6, respectively. Upon cross-referencing with the literature, we found that the outlier 1 Mbp window in chr6 lay in the human leukocyte antigen (HLA), a super-locus responsible for the regulation of the immune system, previously known to be highly polymorphic (Kulski et al., 2022).

The HLA region in the human genome GRCh38 spanned from genomic coordinates 29,602,228 to 33,410,226 on chr6 of the human genome reference assembly GRCh38. Using the ‘Interface’ module, we confirmed from the meta-information of the VCF that the variants were called concerning GRCh38 and proceeded to filter the variants in the HLA region. Summarising the filtered variants, we observed that 29,336 variants were present in the HLA region. We analysed further to identify if these variants were previously known by comparing the filtered VCF file with the 1KGP variants. To this end, we filtered the HLA variants from the entries in chr6 of the 1KGP variants and compared them with the HLA variants in HG002. Of the 4.86M 1KGP variants reported in chr6, 120,482 variants corresponded to the HLA region. From Figure 2F, we observe that 35.6% of the variants called in the HLA region in HG002 were novel to the 1KGP variants. Further analyses using other bioinformatics methods and tools are required to understand the significance of these novel variants.

### SCI-VCF adds features to the existing suite of open-source tools

We compared the main features in SCI-VCF with the existing open-source tools VIVA (Tollefson et al., 2019), vcfR (Knaus and Grünwald, 2017), vcflib (Garrison et al., 2022), CuteVariant (Schutz et al., 2022), re-Searcher (Schutz et al., 2022), VCF-Miner (Hart et al., 2016), VCFtools (Danecek et al., 2011), and GEMINI (Paila et al., 2013). The results are detailed in Table 1. Apart from SCI-VCF, CuteVariant, VCF-Miner, and re-Searcher provided GUIs, while only re-Searcher and GEMINI provided installation-free online access. Visualisation of genetic variants could be handled effectively by only SCI-VCF and VIVA. SCI-VCF is the only tool equipped with a GUI that can summarise, compare and visualise genetic variants in a single platform. We have designed SCI-VCF to make genomic data analyses more open and accessible to all researchers, irrespective of their programming expertise, by simplifying the essential components for VCF-based genetic variant studies. In tasks like sample filtering and variant annotations where SCI-VCF cannot perform optimally, other existing tools execute well, and the resulting VCF files are back-compatible, making their analysis within SCI-VCF feasible. So, SCI-VCF is a valuable augmentation to the current suite of tools rather than a substitute. It complements and enhances existing tools, making genomics more accessible and beneficial for non-programmers.

**Table 1:** Comparison of features of VCF analysis tools.

| Features | SCI-VCF | VIVA | vcflib | vcfR | CuteVariant | re-Searcher | VCF-Miner | VCFtools | GEMINI |
| --- | --- | --- | --- | --- | --- | --- | --- | --- | --- |
| **Technical** |  |  |  |  |  |  |  |  |  |
| **Language** | R | Julia | C++ | R | Python | Python | Java | C++, Perl | Python |
| **Environment (OS)** | Windows, Mac, Linux | Windows, Mac, Linux | Windows, Mac, Linux | Windows, Mac, Linux | Windows, Mac, Linux | Windows, Mac, Linux | Windows | Windows, Mac, Linux | Windows, Mac, Linux |
| **Graphical User Interface** | Yes | No | No | No | Yes | Yes | Yes | No | No |
| **Online version** | Yes | No | No | No | No | Yes | No | No | Yes |
| **Summarise** |  |  |  |  |  |  |  |  |  |
| **Variant level summary** | Yes | No | Yes | Yes* | No | No | No | Yes | No |
| **Sample level summary** | No | Yes | Yes | Yes* | No | No | No | Yes* | No |
| **Compare** |  |  |  |  |  |  |  |  |  |
| **Variant level comparison** | Yes | No | Yes | Yes* | No | No | No | Yes | No |
| **Unique/intersection set summary** | Yes | No | Yes* | Yes* | No | No | No | Yes* | No |
| **Inspect** |  |  |  |  |  |  |  |  |  |
| **Variant-based filtering** | **Yes** | Yes | Yes | Yes | Yes | Yes | Yes | Yes | Yes |
| **Sample-based filtering** | No | Yes | Yes | Yes | Yes | Yes | Yes | Yes | Yes |
| **Visualise** |  |  |  |  |  |  |  |  |  |
| **Interactive plots** | **Yes** | Yes | No | Yes* | No | No | No | No | No |
| **Visualise Genomic regions** | **Yes** | Yes | No | Yes* | No | No | No | No | No |
| **Output** |  |  |  |  |  |  |  |  |  |
| **Publication quality graphics** | **Yes** | Yes | Yes | Yes* | No | No | No | No | No |
| **Export results as tabular data** | **Yes** | Yes | Yes* | Yes* | Yes | Yes* | Yes | Yes | Yes |
| * Multi-step process involving other tools/libraries |  |  |  |  |  |  |  |  |  |

## CONCLUSION

We have developed SCI-VCF as a cross-platform application to summarise, compare, inspect and visualise the genetic variants from the widely accepted variant call format. Researchers and clinicians can use the guided GUI setting of SCI-VCF to perform exploratory genomic data analysis on VCF files, irrespective of their programming expertise. The user-friendly and intuitive design of SCI-VCF will increase the approachability of genomics to newcomers and introduce genomic data analysis expeditiously. We show a case study to illustrate the utility of SCI-VCF. While the use cases pertained to the human genome, the tool is not specific and can be generalised to variants in any organism. We also compared the features in SCI-VCF with other existing VCF analysis software.

Ongoing developments are in progress to enhance the utility of the current version of SCI-VCF. While SCI-VCF offers a wide range of functionalities, we seek to refine its capacity for advanced variant annotation analyses and allele frequency-based filtering. Despite its capability to seamlessly handle VCF files with multiple samples, SCI-VCF does not incorporate a merge function to combine variants from multiple samples. These advanced tasks are resource-intensive, and their incorporation would make maintaining the online version of SCI-VCF expensive. In future versions of SCI-VCF, we aim to expand the framework of SCI-VCF to support the resource-efficient pre-processing of VCF files to add these advanced features successfully. We also intend to parse VCF files parallelly instead of sequentially to handle exceptionally large VCF files efficiently.

## SOFTWARE AND CODE AVAILABILITY

The versatility of SCI-VCF makes it suitable for local installation, allowing personal use and server deployment permitting communal use. SCI-VCF can be used without any installation as an online tool at https://ibse.shinyapps.io/sci-vcf-online/. It can also be installed in diverse ways per the user’s requirements. With R installed, local installation of SCI-VCF is practicable as the dependencies are downloaded automatically upon first use. To work with larger VCF files, it is preferable to use a local installation, as the online version contains resource constraints.

For better package management, a Conda virtual environment with the dependencies of SCI-VCF was created, which can be reproduced easily for improved environment handling. A containerised form of the application is available as a Docker image that helps run SCI-VCF irrespective of platforms, making it straightforward to port the tool to remote HPC clusters. Users can download and install SCI-VCF using the code available on GitHub (https://github.com/HimanshuLab/SCI-VCF). A well-documented guide is available at (https://himanshulab.github.io/SCI-VCF-docs/) for improved user support and system understanding.

## SUPPLEMENTARY INFORMATION

All the analysis in the case study section of the manuscript was done using a system with an M1 processor and 8GB of RAM. The VCF files used in this section are available at https://doi.org/10.5281/zenodo.10780916. The variants were called in the GIAB HG002 sample using the BWA-GATK pipeline containerised at https://hub.docker.com/r/ibse/genome-india.

## ACKNOWLEDGEMENTS

We acknowledge Ayam Gupta for creating the VCF file used in the case study section. We thank Veerendra Gadekar and Preetha Ravi for suggesting features in the tool and proofreading the manuscript. We are grateful to the Centre for Integrative Biology and Systems Medicine (IBSE) and Robert Bosch Centre for Data Science and Artificial Intelligence (RBCDSAI) members for discussions and comments.

